# Noninvasive cancer detection by extracting and integrating multi-modal data from whole-methylome sequencing of plasma cell-free DNA

**DOI:** 10.1101/2022.07.04.498641

**Authors:** Fenglong Bie, Zhijie Wang, Yulong Li, Yuanyuan Hong, Tiancheng Han, Fang Lv, Shunli Yang, Suxing Li, Xi Li, Peiyao Nie, Ruochuan Zang, Moyan Zhang, Peng Song, Feiyue Feng, Wei Guo, Jianchun Duan, Guangyu Bai, Yuan Li, Qilin Huai, Bolun Zhou, Yu Huang, Weizhi Chen, Fengwei Tan, Shugeng Gao

## Abstract

Plasma cell-free DNA (cfDNA) methylation and fragmentation signatures have been shown to be valid biomarkers for blood-based cancer detection. However, conventional methylation sequencing assays are inapplicable for fragmentomic profiling due to bisulfite-induced DNA damage. Here using enzymatic conversion-based low-pass whole-methylome sequencing (WMS), we developed a novel approach to comprehensively interrogate the genome-wide plasma methylation, fragmentation, and copy number profiles for sensitive and noninvasive multi-cancer detection. With plasma WMS data from a clinical cohort comprising 497 healthy controls and 780 patients with both early- and advanced-stage cancers of the breast, colorectum, esophagus, stomach, liver, lung, or pancreas, genomic features including methylation, fragmentation size, copy number alteration, and fragment end motif were extracted individually and subsequently integrated to develop an ensemble cancer classifier, called THEMIS, using machine learning algorithms. THEMIS outperformed individual biomarkers for differentiating cancer patients of all seven types from healthy individuals and achieved a combined area under the curve value of 0.971 in the independent test cohort, translating to a sensitivity of 86% and early-stage (I and II) sensitivity of 77% at 99% specificity. In addition, we built a cancer signal origin classifier with true-positive cancer samples at 100% specificity based on methylation and fragmentation profiling of tissue-specific accessible regulatory elements, which localized cancer-like signal to a limited number of clinically informative sites with 66% accuracy. Overall, this proof-of-concept work demonstrates the feasibility of extracting and integrating multi-modal biomarkers from a single WMS run for noninvasive detection and localization of common cancers across stages.

## Introduction

Cancer is becoming the most deadly disease and responsible for almost 10 million deaths around the globe in 2020 (1). Detecting cancer early provides an opportunity for more effective therapeutic intervention, which may reduce treatment morbidity and mortality (2). In recent years, cfDNA-based liquid biopsy has gained prominent interest in cancer detection and diagnosis with its minimal invasiveness, high sensitivity, and the potential for simultaneous multi-cancer detection and localization. However, the concentration of tumor-originated cfDNA or circulating tumor DNA (ctDNA) is very low especially among early-stage patients (3, 4), calling for the advancement of experimental and computational approaches to improve the signal-to-noise ratio for more accurate cancer detection.

Although cancer genome is characterized with accumulating mutations (5), the amount of mutation-bearing fragments released into circulation is generally low (6, 7). In addition, there is increasing evidence suggesting the presence of somatic mutations in non-malignant tissues (8, 9). These limitations hamper the clinical utilization of mutation search for cancer detection in terms of both sensitivity and specificity. To solve these problems, epigenetic biomarkers are under active investigation in the liquid biopsy field. Epigenetic dysregulation is an early event during carcinogenesis and involves widespread alterations in DNA methylation and chromatin organization across the genome (10, 11). Profiling of the plasma epigenome is therefore expected to improve detection performance over individual point mutations. Meanwhile, epigenetic signatures are cell- and tissue-specific and can provide clues about the origin of cancer signals (12–16).

Existing studies have shown the promise of targeted methylation sequencing of feature CpG sites as an effective approach for single- or multi-cancer detection (14,17–19). However, this method involves complex procedures such as panel design and target capture. We sought to simplify the workflow and reasoned that tumor-associated whole-genome alterations may serve as alternative biomarkers. For example, large-scale methylation changes are characteristic of most types of tumors (20–22) and analysis of hypomethylated segments of the plasma genome shows diagnostic value for hepatocellular carcinoma (23). Similarly, fragment sizes of plasma cfDNA among cancer patients show more variability than healthy individuals and exhibit position-specific patterns, allowing for the employment of genome-wide fragmentation profiling for multi-cancer detection (15,24–26). Another fragmentomic marker is the composition of the 4-mer fragment end motifs whose frequencies are revealed to be perturbed in cancer patients (27). In addition, copy number alteration (CNA) of large chromosomal segments can also be observed in plasma genome and is considered a highly specific feature of cancer (13,28–30).

We reasoned that integrating above genetic and epigenetic biomarkers could achieve more robust performance than profiling individual features alone. However, conventional bisulfite-based methylation sequencing leads to severe DNA damage (31) and a separate whole-genome sequencing (WGS) library would be needed at the expense of more cfDNA materials if concurrent fragmentomic analysis is desired. Recently, a novel methylation sequencing method utilizing enzymatic conversion of cytosine was developed to minimize DNA damage (32). We employed this method and performed whole-methylome sequencing (WMS) of plasma cfDNA and demonstrated the first proof of concept that WMS data is feasible for simultaneous plasma methylomic and fragmentomic profiling. With low-pass WMS data of plasma samples from the multicenter, case-control, observational MONITOR (<u>M</u>ulti-<u>O</u>mics <u>N</u>oninvasive Inspection of <u>T</u>um<u>O</u>r <u>R</u>isk) study, we were able to develop a machine learning classifier termed “<u>TH</u>orough <u>E</u>pigenetic <u>M</u>arker Integration <u>S</u>olution” (THEMIS) to incorporate genome-wide methylation, fragmentation, and CNA features for sensitive detection of seven common cancers. A cancer signal origin (CSO) classifier was also developed based on methylation and fragmentation profiles at tissue-specific accessible regulatory elements. Together, this proof-of-concept study provides a novel approach for sensitive and accurate multi-cancer detection and localization with liquid biopsy.

## Methods

### Study design and participants

Plasma samples were prospectively collected from 497 healthy donors and 780 patients with breast, colorectal, esophageal, gastric, liver, lung, or pancreatic cancers of various stages and analyzed retrospectively. Participants were enrolled from six hospitals including the Cancer Hospital Chinese Academy of Medical Sciences, and protocols were approved by the ethics committee of all participating institutes (NCC– 007821). Written informed consent for research use was provided to all participants prior to enrollment in the study. Individuals were considered healthy if they had no previous history of cancer and negative clinical diagnosis at the time of administration. Plasma samples from cancer patients were obtained before tumor resection or therapy. Participants were randomly assigned into a training cohort and a test cohort at a ratio of 7:3. Clinical information of participants including age, gender, and disease stage (TNM) were summarized in Table S1. Patients with unknown disease stage were indicated as stage NA.

### Isolation of plasma cfDNA

Plasma samples were isolated from 10 ml of peripheral blood collected and stored in Cell-Free DNA Storage Tube (Cwbiotech). Blood was centrifuged at 1,600 g for 10 min at 4 °C and plasma was transferred to a new tube. A second centrifuge was performed at 12,000 rpm for 15 min at 4 °C to remove remaining cell debris and ∼4 ml of plasma was obtained and stored at −80 °C until use. cfDNA was extracted using MagMAX Cell-Free DNA Isolation Kit (Thermo Fisher Scientific) per manufacturer instructions. The quantity and quality of extracted cfDNA was assessed with Bioanalyzer 2100 (Agilent).

### Whole-methylome sequencing of plasma cfDNA

A range of 5 to 30 ng of cfDNA was used to generate WMS libraries with NEBNext Enzymatic Methyl-seq Kit (New England Biolabs) per manufacturer instructions.

Libraries were amplified with 9 cycles of PCR reactions and quantified using Qubit dsDNA HS Assay Kit (Thermo Fisher Scientific). Libraries were sequenced on NovaSeq 6000 (Illumina) with read length of paired-end 100 bp.

### Methylation sequencing data processing

Methylation sequencing reads were demultiplexed by Illumina bcl2fastq and adaptors were trimmed by Trimmomatic (v.0.36). Reads were aligned against the human reference genome (hg19) and deduplicated by BisMark (33). Samtools (v.1.3) and BamUtil (1.0.14) (https://github.com/statgen/bamUtil) were used for sorting and overlap-clipping of mapped reads. Reads with mapping quality below 20 were filtered out. To normalize for sequencing depth among samples, 60 million paired reads were randomly selected from each sample and used for downstream analyses.

### Copy number profiling

Hg19 autosomes were divided into adjacent, non-overlapping 100-kb bins and the coverage of each bin was calculated. A LOESS-based method (13) was applied to correct for coverage bias related with GC content of the reference genome. Bins overlapping the Duke blacklisted regions (http://hgdownload.cse.ucsc.edu/goldenpath/hg19/encodeDCC/wgEncodeMapability/) or the hg19 gap track (downloaded from UCSC Table Browser) were prone to low mapping quality and removed from analysis. Bins with original coverage over 100 but zero coverage after GC-correction were also excluded. To remove bins with copy number variations among healthy controls, we calculated the z-score of each GC-corrected bin over the 352 healthy controls in the training cohort (i.e., baseline samples). Bins with absolute z-scores above two among more than 60 baseline samples or above four among any baseline samples were filtered out.

### Fragment size index (FSI)

To profile differences in cfDNA fragmentation size between cancer patients and healthy controls, the frequency of each fragment size was calculated for individual samples and Wilcoxon test was performed between cancer samples and healthy controls in the training cohort to select for differentially represented fragment sizes (Figure S1). At a p value threshold of 0.01, most fragment sizes within 50–300 bp exhibit frequency differences except 167–168 bp and 241–259 bp. Therefore, fragments within 100–166 bp were considered as short fragments and fragments within 169–240 bp were considered as long fragments. The ratio of short to long fragment counts was defined as fragment size index (FSI).

To characterize the genome-wide FSI profile, hg19 autosomes were divided into 100-kb A/B compartments representing open and closed chromatin regions (34) and those overlapping the Duke blacklisted regions or the hg19 gap track were removed from analysis. Counts of short and long fragments within each bin was calculated and the ratios of short to long fragment counts were corrected against GC content of the reference genome using a LOESS-based method (13). After merging adjacent 100-kb bins, genome was segmented into 502 non-overlapping 5-Mb windows (Table S2) and the FSI of each 5-Mb window was defined as the average of its overlapping GC-corrected 100-kb FSI values.

### Methylated fragment ratio (MFR)

Hg19 autosomes were tiled into 1,846 adjacent, non-overlapping 1-Mb windows after filtering out genomic regions overlapping the Duke blacklisted regions or the hg19 gap track (Table S3). Read pairs were merged into fragments and those failing to meet the following criteria were discarded: (1) Fragment covers at least 3 CpGs. (2) Fragment length is between 80 and 250 bp, which is the size range of most cfDNA molecules. (3) Conversion rate of non-CpG cytosines exceeds 95%. The methylation level of each 1-Mb window was quantified by the fraction of fragments with fully methylated CpGs, or methylated fragment ratio (MFR).

### Associations among fragmentation, methylation, and copy number profiles

The 1,846 1-Mb windows determined by MFR analysis were used to investigate the associations among methylation, fragmentation, and CNA across the genome. The FSI of each window was calculated as the average GC-corrected FSI values of its overlapping 100-kb bins. For CNA analysis, the coverage of each window was calculated as the total GC-corrected coverage of its overlapping 100-kb bins. Because of the more stringent filtering criteria for CNA analysis, 1,795 windows were eventually kept for association studies. Each feature type for a sample was quantile-normalized with that of healthy controls. A z-score was calculated for each bin over the corresponding bins of healthy controls.

### Chromosomal aneuploidy of featured fragments (CAFF)

CAFF analysis followed the same procedures as CNA profiling except that fragments shorter than 151 bp or longer than 220 bp were extracted for coverage analysis to increase the presence of tumor-derived abnormal fragments (35, 36). The coverage of each chromosome arm was calculated by summing the GC-corrected coverage of all its 100-kb bins. To summarize chromosome arm-level copy number changes, we adopted a previously described approach to calculate the plasma aneuploidy (PA) score of each sample using the five chromosome arms exhibiting the most dramatic copy number alterations from baseline samples (13).

### Fragment end motif (FEM)

The 4-nucleotide (i.e., 4-mer) fragment 5’ end motif was extracted as previously reported (27) with the following modifications: (1) Fragments shorter than 171 bp were selected for analysis. (2) Only reads mapped to the Crick strand were used for calculation.

### Machine learning models for MFR, FSI, and FEM

An in-house Python script based on sklearn module was used for the development of machine learning models. To mitigate overfitting, 13 bootstraps of the training cohort was performed for hyperparameter optimization and the average probability of the resulting 13 sub-models was used as the prediction score of a sample. Model performance for each biomarker was evaluated with the independent test cohort.

#### MFR

To construct an MFR classifier, principal component analysis (PCA) was performed for the MFR values of the 1,846 windows to reduce feature dimensionality. The first 53 principal components (PCs), which explained at least 95% of the variance of the training cohort, were used for model development. A support vector machine (SVM) model was trained with the training cohort under 10-fold cross-validation.

#### FSI

For each sample, the FSI values of the 502 5-Mb windows were normalized to z-scores across the genome. PCA was performed and the top 42 PCs which explained at least 90% of the variance of the training cohort was used to develop an SVM model under 10-fold cross-validation.

#### FEM

The frequencies of the 256 4-mer end motifs of each sample were quantile-normalized with the healthy controls in the training cohort. PCA was performed and the top 31 PCs explaining at least 90% of the variance of the training cohort was used to develop a logistic regression (LR) model under 10-fold cross-validation.

### Development of THEMIS classifier for biomarker integration

To integrate the predicted cancer probability scores by MFR FSI, and FEM along with the PA scores of CAFF, a generalized linear model (GLM) with elastic-net penalization was constructed with the R package CARET under 20-fold cross validation. The best tuned elastic-net mixing parameter alpha is 0.079. The resulting THEMIS model calculates the probability of cancer in a participant as:

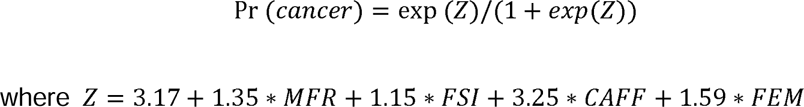

Prediction scores by individual biomarkers and the integrative THEMIS classifier were listed in Table S4.

### Cancer signal origin classification

We chose cancer samples identified as true positives by THEMIS at 100% specificity to develop and evaluate the CSO classifier. Plasma methylation and fragmentation profiles for 18 clusters of tissue-specific 500-bp open chromatin peaks identified by ATAC-seq (22) were used for model development.

For each sample, methylation level was quantified by the fraction of methylated CpGs within individual ATAC-seq peaks. Because open chromatin regions tend to be hypomethylated (22, 39), Wilcoxon test was performed between cancer samples and healthy controls in the training cohort to select for peaks with lower methylation among cancer patients (p < 0.05), which constituted ∼90% of the original ATAC-seq peaks, for CSO analysis. For each cluster, the average methylation level of the remaining peaks was computed and transformed into a z-score over the corresponding cluster of healthy controls in the training cohort. Subsequently, the z-scores of 18 clusters were scaled between 0 and 1 for each sample.

To characterize the fragmentation profile of each sample, the aggregate counts of short (100–166) and long (169–240 bp) fragments falling within the peak regions of each cluster were calculated respectively and transformed into z-scores over the corresponding cluster of healthy controls in the training cohort. The z-scores of 18 clusters were then scaled between 0 and 1 for each sample.

Finally, the normalized methylation level, short fragment coverage, and long fragment coverage of each cluster were used as the input features for the development of the multi-class CSO classifier. A random forest model was trained using RandomForest package in R with 2,000 trees under leave-one-out cross-validation.

## Results

### Overview of THEMIS workflow

The workflow of THEMIS involves the development of both experimental and computational platforms (Figure 1). To generate plasma WMS libraries, we isolated cfDNA from ∼4 mL of plasma and applied a range of 5–30 ng of cfDNA as input materials. Enzymatic reactions, including protection of 5-mC and 5-hmC by TET2 (Tet Methylcytosine Dioxygenase 2) and subsequent cytosine deamination by APOBEC (Apolipoprotein B mRNA Editing Catalytic Polypeptide-like), were utilized to convert unmodified cytosine into uracil (32). WMS libraries were subject to low-pass paired-end sequencing, and all uniquely aligned sequencing data were randomly down-sampled to 60 million (M) paired reads (∼2X haploid genome coverage) for downstream analysis.

**Figure 1.**
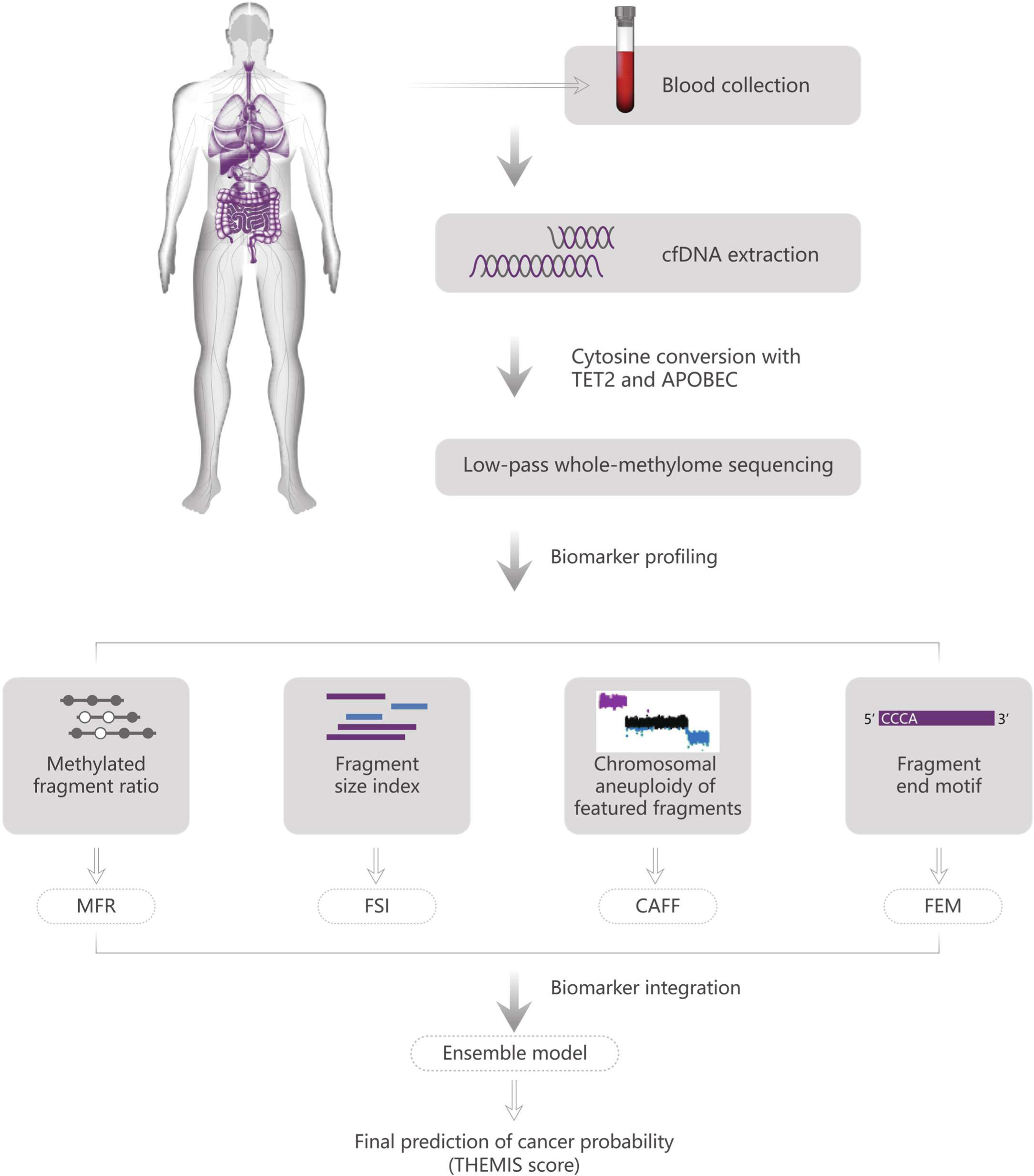
Overview of THEMIS approach for cancer detection based on plasma whole-methylome sequencing. Schematic illustration of the experimental and bioinformatics procedures. Blood samples were collected from cancer patients or healthy donors. Plasma cfDNA was extracted from the participant’s blood sample and subject to low-pass WMS using TET2 and APOBEC enzymes for cytosine conversion. Four types of biomarkers were extracted from uniquely mapped WMS sequencing reads, including Methylated Fragment Ratio (MFR), Fragment Size Index (FSI), Chromosomal Aneuploidy of Featured Fragments (CAFF), and Fragment End Motif (FEM). An ensemble model integrating prediction scores from all four biomarkers was constructed to yield the final probability of having cancer, termed “THEMIS score”, for a sample.

Given that the mild enzymatic reactions minimize DNA damage, we sought to incorporate CNA and fragmentomic analyses besides methylation profiling for more sensitive cancer detection. We designed algorithms to extract four genetic and epigenetic cancer biomarkers from plasma cfDNA. To profile the genome-wide methylation patterns, we divided the genome into 1,846 non-overlapping 1-Mb segments and calculated the ratio of fully methylated fragments within each window (“Methylated Fragment Ratio”, MFR). Similarly, position-specific fragmentation characteristics were profiled as the ratios of short (100–166 bp) to long (169–240 bp) fragments for 502 non-overlapping 5-Mb genomic windows (“Fragment Size Index”, FSI). To enhance the signal for copy number analysis, we size-selected short (< 151 bp) and long (> 220 bp) fragments which are more likely to originate from cancer cells (35, 36) to quantify the copy number changes of chromosome arms (“Chromosomal Aneuploidy of Featured Fragments”, CAFF). In addition, the frequencies of 256 4-mer motif patterns at the 5’ end of fragments were quantified (“fragment end motif”, FEM). Machine learning methods were implemented for MFR, FSI, and FEM profiling to improve the performance of cancer detection. Finally, an ensemble classifier was constructed by integrating individual biomarkers to predict a sample’s probability of having cancer, which was termed the THEMIS score.

### WMS is feasible for cfDNA CNA and FSI profiling

The applicability of WMS data for CNA and fragmentomic profiling sets the foundation of THEMIS. To confirm that enzyme-converted WMS libraries can be adapted to characterize these genomic features, we assessed the concordance of CNV and FSI profiles between WMS and the “gold-standard” WGS platforms (13,15,24,26,37) generated from paired plasma samples of the same individuals.

As observed for the plasma genome of a late-stage colorectal cancer patient (GCP0088), WMS displayed highly consistent copy number patterns with WGS in genomic bins of 100 kb as well as aneuploidy of the same chromosomes, including gains of chromosomes 13 and 20 and losses of chromosomes 4 and 18 that frequently occur in colorectal cancer (38) (Figure 2A). We used PA score which summarized copy number changes of the top five chromosome arms (13) to evaluate genome-wide aneuploidy of a sample. Among a clinical cohort comprising 220 healthy controls and 270 cancer patients of multiple types (Table S5), PA scores were closely matched between WMS and WGS data with a Pearson correlation of 0.988 (95% confidence interval (CI): 0.986–0.990) (Figure 2B), suggesting that WMS could achieve comparable performance to WGS platform in plasma CNA detection.

**Figure 2.**
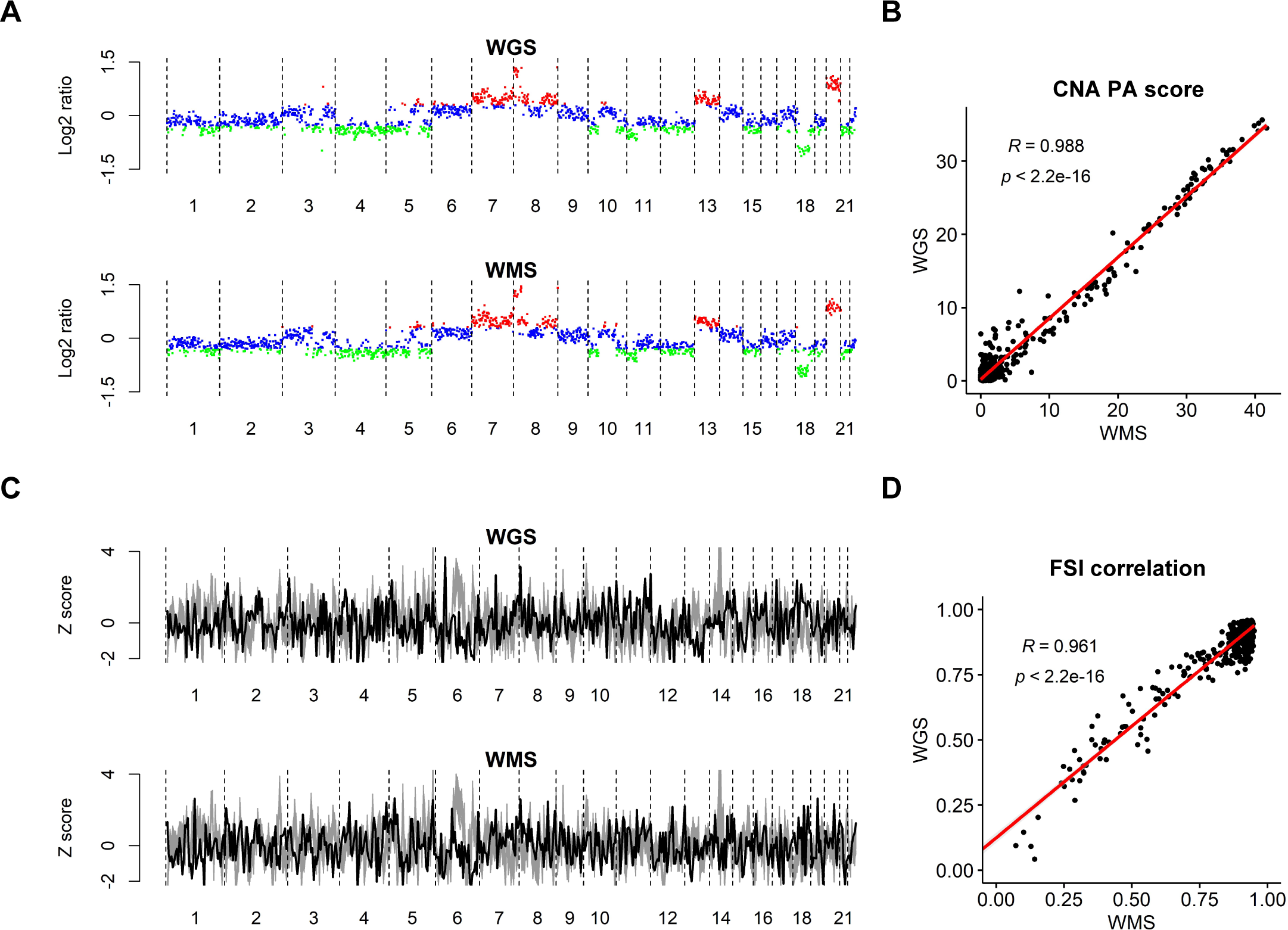
Concordance between WMS and WGS in cfDNA copy number and fragmentation profiling. (A) and (C) depict the genome-wide CNA and FSI profiles of a colorectal cancer patient GCP0088 as an example. (B) and (D) analyze 225 healthy controls and 287 cancer patients for cohort-level comparison. (A) Log2 ratio over the mean baseline coverage in 100-kb bins across the genome of Patient GCP0088 profiled by WGS and WMS respectively. Bins with log2 ratio above 0.3 (copy number gains) are colored in red, and bins with log2 ratio below -0.3 (copy number loss) are colored in green. (B) Scatter plot of PA scores derived from paired WGS and WMS data of all samples. The regression line is colored in red. (C) FSI of 5-Mb windows across the genome of Patient GCP0088 profiled by WGS and WMS respectively. The coverage ratio of short to long fragments for each bin is normalized by z-score across the genome. Patient FSI profile is colored in black against 225 grey healthy reference FSI profiles. (D) Scatter plot of Pearson correlations between FSI profiles of individual samples with that of the median healthy reference, assessed with WGS and WMS data respectively. The regression line is colored in red.

The concordance of cfDNA fragmentation profiles between WMS and WGS data was analyzed in a similar manner. Across the plasma genome of Patient GCP0088, short to long fragment ratios in 5-Mb windows showed similar trends between the two platforms (Figure 2C). With the same cohort as for CNA assessment, we used Pearson correlation to quantify the similarity of a sample’s genome-wide fragmentation profile to that of the healthy reference (i.e., the median ratio of healthy controls for each window). Among all cohort samples, WMS and WGS platforms showed high concordance in FSI profiling with a Pearson correlation of 0.961 (95% CI: 0.954–0.968) (Figure 2D). These data demonstrate that WMS is also feasible for cfDNA fragmentation analysis.

### Associations among methylation, fragmentation, and copy number alterations of the plasma genome

The multi-modal biomarkers we profiled may represent different types of molecular alterations of common ctDNA fragments (22) and therefore are connected with each other. For example, genomic segments with copy number changes tend to exhibit more dramatic fragmentation alterations (15, 24), perhaps corresponding to over- or under-representation of ctDNA fragments shed from these regions. To investigate the associations among fragmentation, methylation, and copy number, we selected 20 healthy controls and 20 colorectal cancer samples (Table S6) and profiled their FSI, MFR, and CNA in 1-Mb windows across the plasma genome. Unlike healthy controls, the genome of cancer patients exhibited both increases and decreases in the signals of all three biomarkers spanning broad regions (Figures 3A to 3C). A large fraction of the altered windows was shared among cancer patients: 10.6% for FSI, 35.5% for MFR, and 40.2% for CNA. In contrast, we identified no commonly altered windows for any biomarker in healthy controls (Figure 3D).

**Figure 3.**
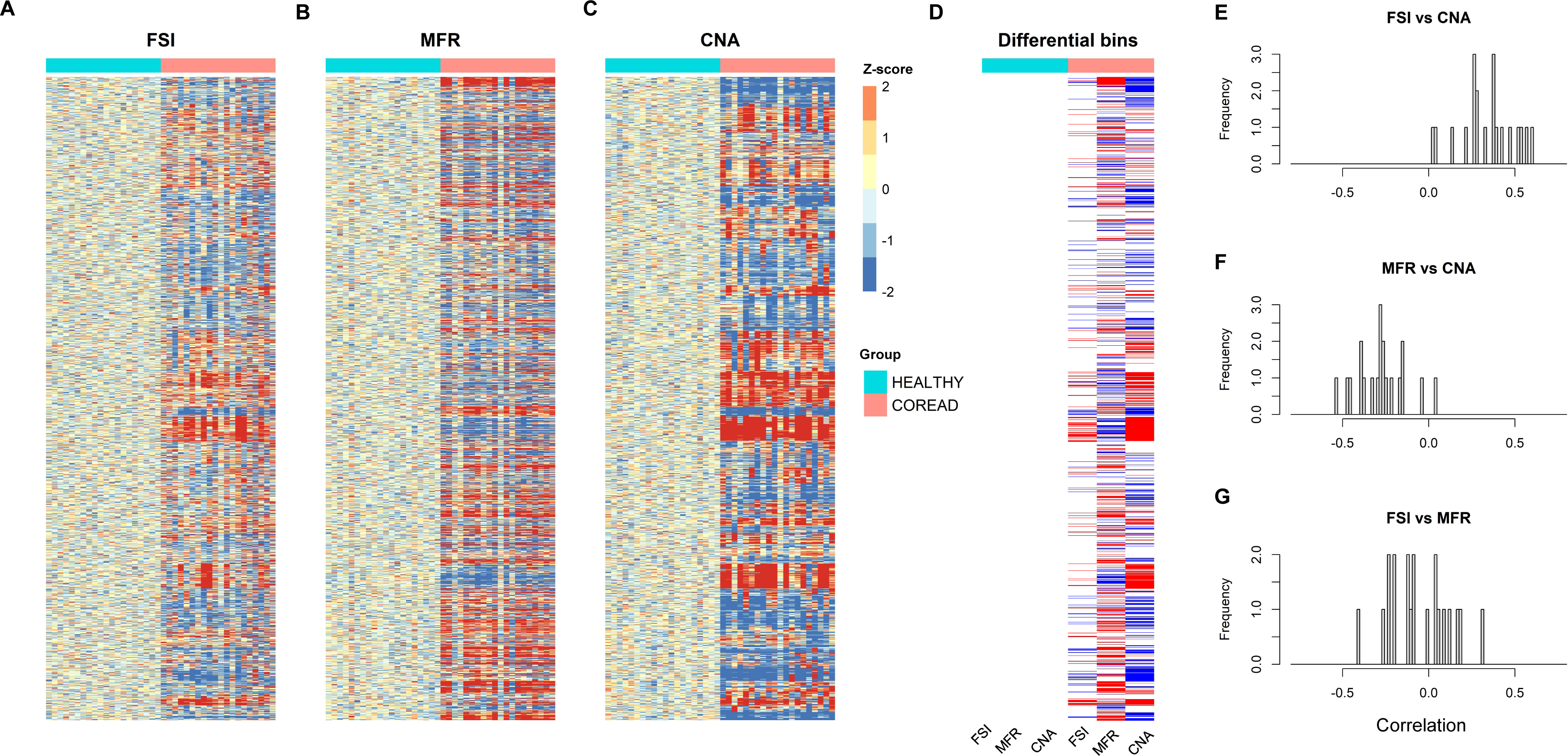
Associations among plasma fragmentation, methylation, and CNA profiles. A total of 20 healthy controls and 20 advanced colorectal cancer patients are analyzed. (A-C) Z-scores of FSI, MFR, and CNA profiles in 1,795 non-overlapping1-Mb windows across the genome (row) of healthy and patient individuals (column). (D) For each feature type, genomic windows are indicated in red if more than half of the healthy controls or more than half of the cancer patients have z-scores above 2 and in blue if z-scores are below −2. (E-G) Distributions of Pearson correlations between the genome-wide FSI and CNA (E), MFR and CNA (F), and FSI and MFR (G) profiles for cancer patients.

To quantify the associations among the three biomarkers, we calculated their pairwise Pearson correlations across all genomic windows for each cancer patient. Consistent with the notion that cancer signals derived from CNV-positive regions are better manifested, positive correlations between FSI and CNA profiles (median r = 0.350) were noted for most patients (Figure 3E). In contrast, MFR and CNA profiles were mostly anti-correlated (median r = −0.276; Figure 3F), probably reflecting widespread hypomethylation of the tumor genome (23). The correlations between FSI and MFR profiles appeared more vague (median r = −0.087; Figure 3G), which might imply mixed relationships between fragmentation size and methylation level. These associations among different types of biomarkers may facilitate in the improvement of detection power if integrative analysis was conducted.

### Biomarker extraction from cfDNA WMS data for cancer detection

To develop and validate the computational approaches for cancer detection by WMS-derived biomarkers, we performed WMS of plasma cfDNA from a case-control clinical cohort comprising 780 previously untreated cancer patients and 497 healthy controls, which were randomly split into a training cohort and an independent test cohort at a ratio of 7:3. The training cohort included 352 healthy controls and 542 cancer patients (46 breast (BRCA), 105 colorectal (COREAD), 42 esophageal (ESCA), 78 liver (LIHC), 110 lung (NSCLC), 83 pancreatic (PACA), and 78 gastric (STAD) cancers), 35.1% of which were at early stages (I or II). The test cohort consisted of 145 healthy controls and 238 cancer patients (20 BRCA, 45 COREAD, 19 ESCA, 35 LIHC, 47 NSCLC, 36 PACA, and 36 STAD), among which 34.5% were at early stages (I or II). We also implemented machine learning methods on individual biomarkers to increase cancer detection performance. It is to be noted that all models were built solely on samples in the training dataset.

The detection performance of individual biomarkers was assessed with receiver operator characteristic (ROC) analysis (Figures 4A). Specifically, to classify whether the methylation landscape (i.e., MFR values of 1, 846 1-Mb windows) of a cfDNA sample is more characteristic of healthy controls or cancer patients, we applied a principal component analysis (PCA) to reduce the dimensionality of the feature space and trained a support vector machine (SVM) model to achieve an area under the ROC curve (AUC) value of 0.946 (95% CI: 0.924–0.968) for the detection of all seven cancer types combined in the test cohort. Similarly, PCA was performed for the FSI profile of the 502 5-Mb windows and an SVM classifier was constructed and achieved an overall testing AUC value of 0.910 (95% CI: 0.882–0.938). CNA analysis of size-filtered fragments by PA scores to summarize chromosomal aneuploidy had a testing AUC value of 0.861 (95% CI: 0.825–0.897). Finally, a logistic regression (LR) model was trained for the 256 4-mer fragment end motifs after PCA and the testing AUC reached 0.932 (95% CI: 0.908–0.956). While individual biomarkers displayed varying detection performances among cancer types, AUC values and prediction scores were consistent between the training and test cohorts for the prediction of individual cancer types (Figures S2-S5, Panels A&B), suggesting minimized overfitting during model development. Moreover, all biomarkers yielded increasing prediction scores with advancing histological stage (Figures S2-S5, Panel C).

**Figure 4.**
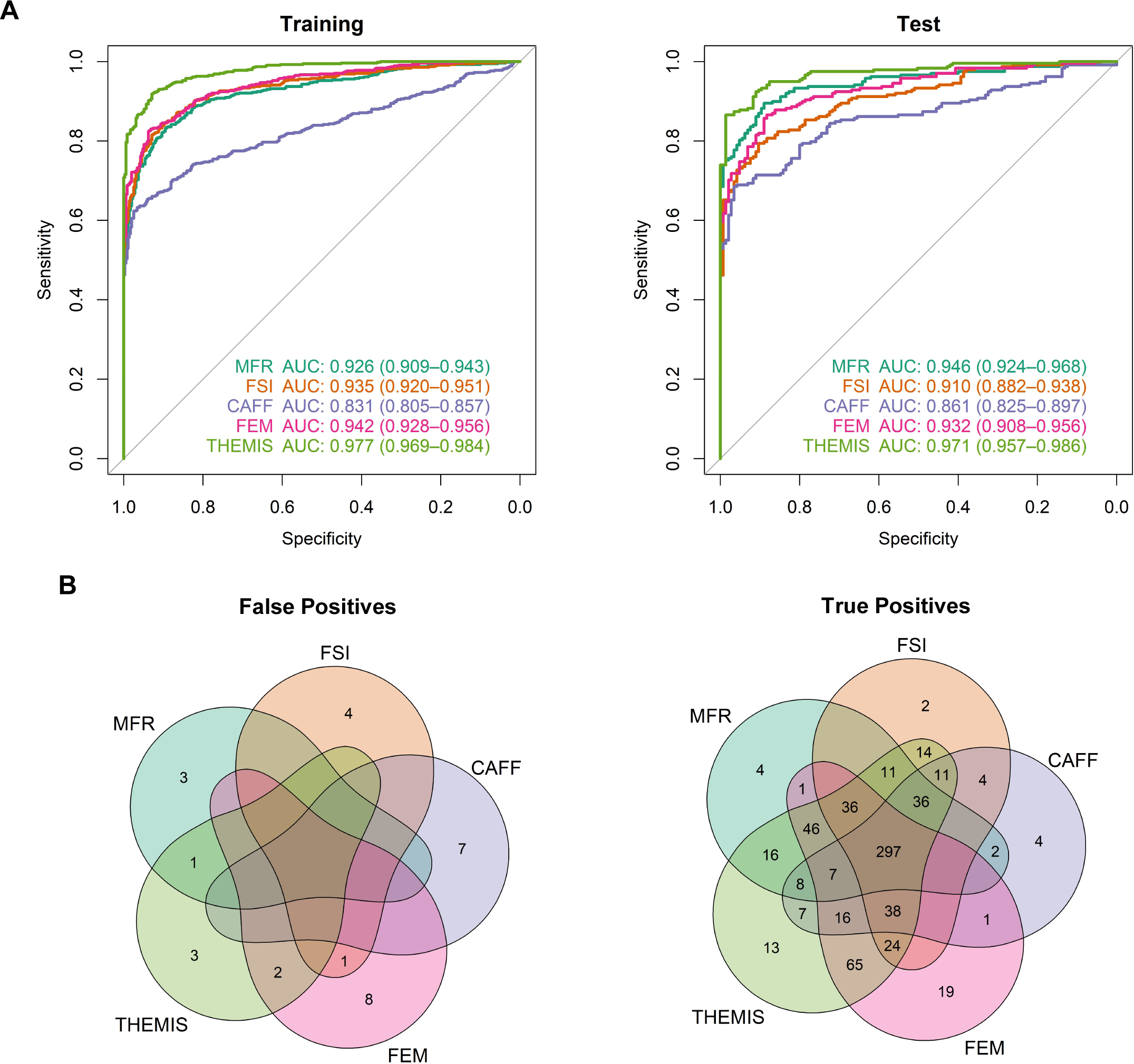
Cancer detection by multi-modal analysis of cfDNA WMS data. (A) Receiver operator characteristics for the classification of cancer patients (training n = 542; test n = 238) from healthy controls (training n = 352; test n = 145) by individual biomarkers and the integrative THEMIS model for the training and the test cohort respectively. (B) Venn diagrams depicting the number of healthy individuals misclassified as cancer patients (false positives) and the number of cancer patients correctly identified (true positives) at a specificity of 99% by individual biomarkers and THEMIS. Samples from both the training and the test cohort are included.

### Integration of biomarkers improves detection power

Prediction scores of individual biomarkers showed strong pairwise correlations but were also distinct for specific samples (Figure S6), suggesting complementary roles of these multi-dimensional biomarkers. We hypothesized that integrative analysis of all four biomarkers could further boost the prediction power and constructed an ensemble THEMIS classifier with a regularized LR model. THEMIS outperformed any single biomarkers with a higher overall AUC value of 0.971 (95% CI: 0.957– in the test cohort (Figure 4A). At a threshold of 99% specificity, THEMIS also reduced the number of healthy individuals misclassified as cancer patients (false positives) and increased the number of cancer patients correctly identified (true positives) (Figure 4B). These results confirm our hypothesis that comprehensive analysis of WMS data can improve both detection specificity and sensitivity over profiling individual feature types alone.

THEMIS achieved high AUC values in the detection of all seven types of cancer ranging from 0.947 (95% CI: 0.911–0.983) for NSCLC to 0.996 (95% CI: 0.990–1.000) for ESCA in the test cohort (Figure 5A). Remarkably, THEMIS maintained consistent AUC values between the training and test cohorts for every cancer type, suggesting minimal overfitting of the ensemble classifier. At a specificity of 99%, THEMIS correctly classified 443/542 (82% sensitivity; 95% CI: 78–85%) cancer patients in the training cohort versus 204/238 (86% sensitivity; 95% CI: 81–90%) in the test cohort (Figures 5B&S7A). The prediction scores and detection sensitivity of THEMIS increased with increasing clinical stages (Figures 5B&S7B), likely implying that cancer-like signals profiled by THEMIS were correlated with tumor load. The overall detection sensitivity of THEMIS for early-stage (I and II) cancers was 73% (95% CI: 66%–78%) and 77% (95% CI: 67–85%) in training and test, respectively.

**Figure 5.**
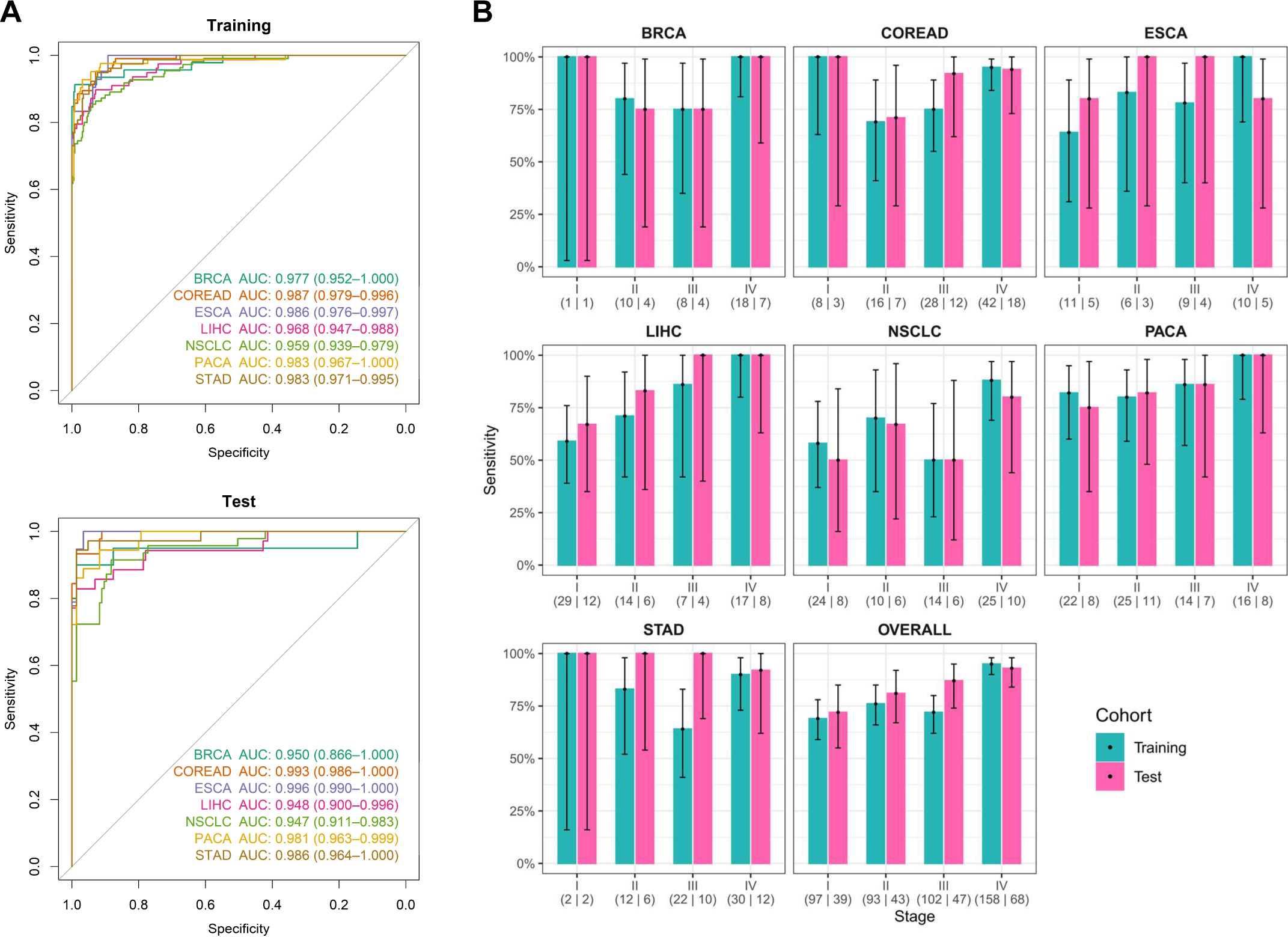
THEMIS performance for multi-cancer detection. (A) Receiver operator characteristics for the classification of cancer patients from healthy controls by THEMIS in the training (healthy n = 352; cancer n = 542) and the test (healthy n = 145; cancer n = 238) cohort split by cancer type. (B) Detection sensitivity of individual cancer types by clinical stage. Sensitivity at a specificity of 99% is depicted with 95% confidence interval for individual cancer types. Clinical stage and the number of samples in the training and the test cohort (separated by a vertical line) are indicated below the plot. Cancer samples with unknown stages are omitted from display.

### Classification of cancer signal origin

Epigenetic signatures of circulating cfDNA may inform the identity of its originating cells (14–16). To locate the origin of cancer signals with low-pass WMS data, we consult to the aggregate methylation and fragmentation profiles at cancer tissue-specific accessible regulatory elements annotated by ATAC-seq (22). We used cancer samples correctly identified as true positives by THEMIS at 100% specificity for CSO analysis, which included 384 training and 176 test samples, and characterized the methylation level and the coverage of short (100–166) and long (169–240 bp) fragments for 18 ATAC-seq clusters in each sample. To orthogonally validate the tissue-specificity annotation of ATAC-seq clusters, we first profiled the normalized signal of individual feature types for each cluster among cancer types with training samples (Figure 6A). Consistent with published reports on the anticorrelation between DNA methylation and chromatin accessibility (22, 39), we observed lower methylation of ATAC-seq clusters in relevant cancer types compared with other cancers (Wilcoxon test). For example, COREAD samples were notably hypomethylated in Cluster 2 (colon) and both COREAD and STAD samples were hypomethylated in Cluster 13 (digestive). Likewise, BRCA and NSCLC samples had lower methylation in Cluster 3 (Non-basal breast) and Cluster 12 (Lung), respectively. We also observed lower coverage of short or long fragments in cancer relevant clusters, which largely resembled methylation profile. Based on the aggregate signals of each cluster for these three feature types, a CSO classifier to differentiate the seven cancer types was developed using a random forest algorithm. This model correctly identified the origin of cancer-like signals for 191/384 (50%, 95% CI: 45%–55%) samples in the training cohort and 90/176 (51%, 95% CI: 44%–58%) samples in the test cohort (Figure 6B). Considering that ESCA, STAD, and COREAD share common ATAC-seq clusters and diagnostic methods in the clinic, we combined these three cancer types into a “digestive cancer” group for CSO assessment. Similarly, we combined LIHC and PACA into the “hepatopancreatic cancer” group. Our CSO classifier achieved an accuracy of 239/384 (62%, 95% CI: 57%–67%) for the training and 116/176 (66%, 95% CI: 59%–73%) for the test samples after cancer type grouping.

**Figure 6.**
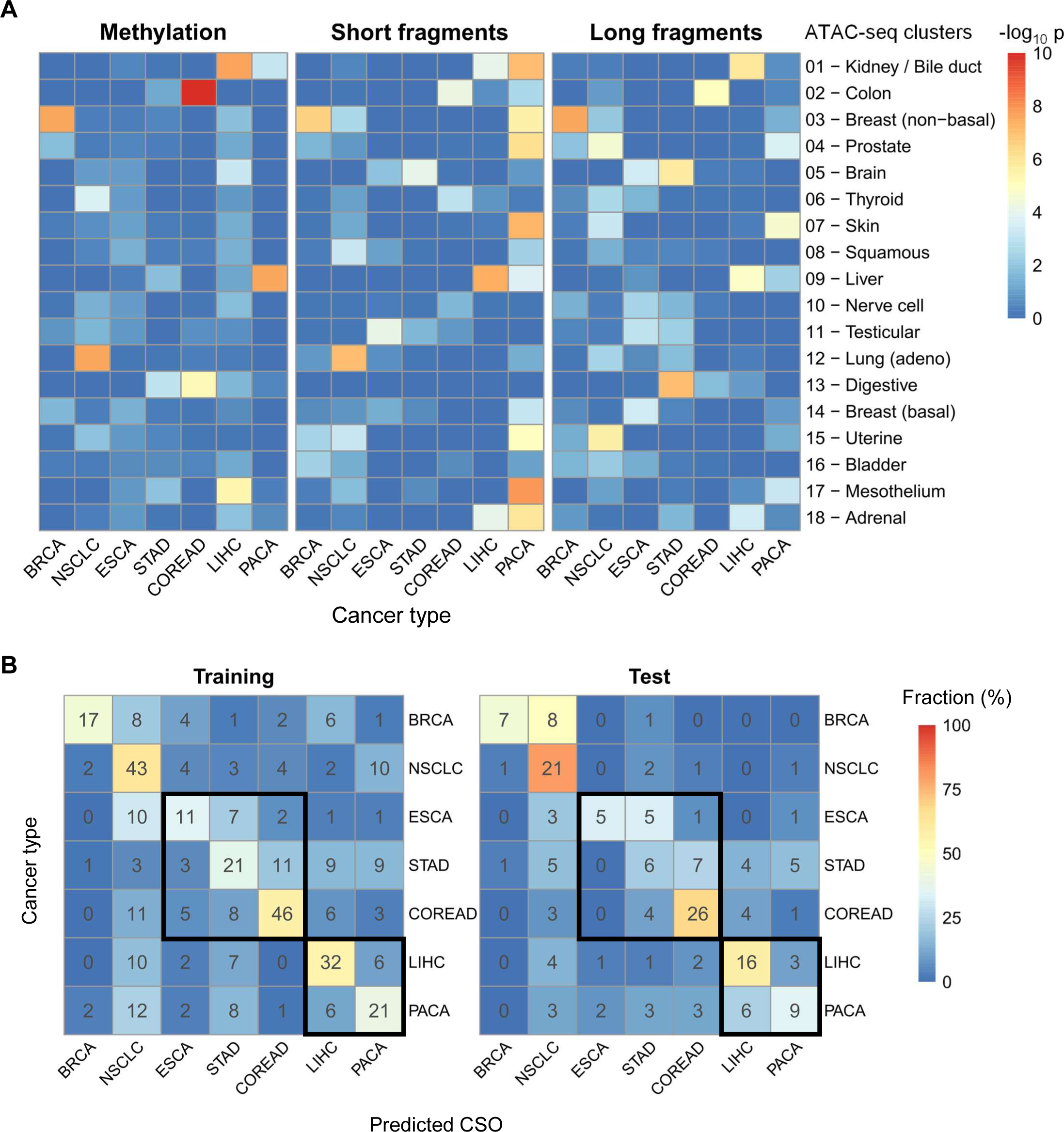
Classification of cancer signal origin. (A) Heat maps of p values (one-sided Wilcoxon test) of methylation levels, short fragment coverage, and long fragment coverage of each ATAC-seq cluster between individual cancer types and other cancers in the training cohort. Each feature is normalized as described in Methods. Training cancer samples correctly identified as true positives by THEMIS score at 100% specificity are used for analysis, including 39 BRCA, 68 NSCLC, 32 ESCA, 57 STAD, 79 COREAD, 57 LIHC, and 52 PACA samples. (B) Confusion matrices representing the accuracy of CSO localization in the training (top) and the test (bottom) cohort. Color corresponds to the proportion of predicted CSO calls. Test cancer samples correctly identified as true positives by THEMIS score at 100% specificity are used for evaluation of the CSO classifier, including 16 BRCA, 26 NSCLC, 15 ESCA, 28 STAD, 38 COREAD, 27 LIHC, and 26 PACA samples. Digestive cancer group and hepatopancreatic cancer group are squared.

## Discussion

In this work, we developed a novel noninvasive multi-cancer detection and localization approach, THEMIS, by leveraging crucial tumor genetic and epigenetic characteristics across the plasma genome with low-pass WMS data. We showed the first proof of principle that enzyme-based WMS platform is applicable for cfDNA fragmentation and coverage analyses, which enables THEMIS to comprehensively profile multiple types of proven ctDNA biomarkers including methylation, fragmentation size, CNA, and end motif patterns. Assessed with plasma samples from a case-control clinical cohort comprising seven cancer types, the integrative THEMIS model outperformed individual biomarkers and demonstrated high sensitivity in cancer detection. Furthermore, THEMIS exhibited great potential in assigning the origin of cancer signals by profiling methylation and fragmentation characteristics of tissue-specific clusters of accessible regulatory elements.

Recently published blood-based multi-cancer detection tests employed targeted methylation sequencing (14), immunoprecipitation-based methylation profiling (16), genome-wide fragmentation patterns (15), or detection of mutation and serum protein biomarkers (40) for cancer detection and classification. Like these multi-cancer detection approaches, THEMIS presents several major advantages over current recommended clinical diagnostic methods such as low-does computed tomography for lung cancer (41) and endoscope for esophageal cancer (42): (1) contemporary imaging technologies require sufficient tumor size for visibility; this limitation can be overcome by blood-based THEMIS test to detect smaller (i.e., earlier) cancers with better sensitivity and accuracy. (2) THEMIS enables multi-cancer detection and CSO localization from a single assay, which avoids the accumulation of false positive rates if multiple individual tests were conducted. (3) THEMIS involves minimally invasive biopsy procedures and is free of radiation exposure, therefore improved patient compliance is anticipated. Moreover, although previous studies have explored integrative analysis of methylation with fragmentation or CNA for better detection sensitivity, methods used therein rely on bisulfite conversion or antibody immunoprecipitation for methylation sequencing and a separate WGS library is required for parallel fragmentation or coverage analysis (23, 37). We showed that WMS platform is comparable to the “gold-standard” WGS platform in cfDNA fragmentation and CNA analyses (Figure 2). The capability of simultaneous profiling of plasma methylome, fragmentome, and copy number with a single WMS library prepared from only 10 mL of whole blood makes THEMIS applicable for low-input samples with substantially simplified experimental workflow. In addition, THEMIS was developed on low-pass sequencing with 60M paired reads. We further subsampled WMS data of select healthy and cancer samples with 60M reads to 45M, 30M, 15M, and 3M reads and noticed highly consistent genome-wide profiles of all individual biomarkers extracted from 30M data with the original 60M data (Figure S7). Therefore, THEMIS can potentially achieve comparable predictive power at even lower sequencing depth thus reducing sequencing cost. Together, these clinical and technical benefits highlight the potential of THEMIS to be employed as a routine population-level cancer screening test.

The concentration of circulating tumor DNA is generally low especially among patients with early-stage disease. To capture weak tumor signals, machine learning methods are frequently implemented to increase the sensitivity of detection which are prone to overfitting issues arising from technical and biological noises. Multiple lines of evidence suggest that THEMIS characterizes bona fide tumor-derived signals. First, because alteration of genome copy number is a highly cancer-specific event (13,28–30), CNA may be utilized to infer the fidelity of cancer signals we detected.

Indeed, across the genome of plasma samples from example colorectal cancer patients, we observed pronounced correlation between FSI and copy number in CNA-positive genomic regions in agreement with other reports (15,24,25); meanwhile we noticed strong anti-correlation between methylation and copy number in these regions (Figure 3). Second, we noticed consistent cancer detection performance between the training and the test cohort with both the base models of individual biomarkers and the integrative THEMIS model (Figure 4), suggesting mitigated overfitting issues if any during model development. Finally, prediction scores and detection sensitivity of THEMIS increased with increasing disease stage (Figures 5&S7), implying that THEMIS signal originates from tumor cells instead of background noise.

Localizing the origin of cancer-like signal is a crucial component of blood-based multi-cancer detection test to provide guidance on the follow-up diagnostic workup. Tissue specificity has been reported for plasma epigenetic features including methylation, fragmentation, and nucleosome footprints, all of which hold promise for the tracing of tumor locations (14,15,43). Among these exploratory studies, high-depth targeted sequencing of signature methylation sites showed robust CSO classification performance (14). Because of the low sequencing depth of WMS data, THEMIS adopted an alternative CSO analysis strategy combining methylation and fragmentation profiles of ATAC-seq data-derived cancer tissue-specific clusters of accessible regulatory elements (22). As expected, we observed hypomethylation and low coverage of cancer type-relevant clusters (e.g., Cluster 2 for COREAD and Cluster 13 for COREAD and STAD) with plasma WMS data, confirming the concordance between tissue and plasma tumor DNA as well as the fidelity of methylation and fragmentation signals extracted from WMS data. Surprisingly with ATAC-seq Cluster 9 (liver) in our LIHC samples, we did not observe strong hypomethylation but the coverage of short and long fragments was significantly reduced. This might reflect histological heterogeneity between cancer samples analyzed in the current study and the original ATAC-seq study (22), as well as complementary roles between methylation and fragmentation profiles in CSO classification. Given the constraining availability of chromatin accessibility datasets, CSO performance presented in this study is a proof of concept for the feasibility of low-pass WMS data to perform this analysis, and we combined cancer types with similar ATAC-seq cluster features during the assessment of CSO accuracy since the grouped cancers share identical clinical diagnostic procedures (gastrointestinal endoscope for digestive cancer and abdominal ultrasound scan for hepatopancreatic cancer, respectively). We expect that additions of chromatin accessibility datasets will expand the types of cancer classifiable by THEMIS and increase model performance.

Despite the proof of concept for a novel sensitive and accurate multi-cancer detection and localization approach, this study has limitations. Due to the small sample size, participant demographics are unmatched between cancer patients and healthy controls. We also lack complete one-year follow-up information for participants considered healthy at the time of writing this manuscript, therefore we cannot rule out the possibility that some control individuals bear early-stage cancer and may overestimate the false positive rates of THEMIS. Better assessment of real-world THEMIS performance and the establishment of its clinical utility require future investigations in a larger cohort comprising age- and sex-matched high-risk healthy controls as well as type- and stage-balanced cancer patients. Complete long-term follow-up is also necessary.

In conclusion, this proof-of-concept study demonstrates the robust detection performance and CSO classification feasibility of common cancers for WMS-based THEMIS approach. These results support THEMIS as a novel affordable population-level multi-cancer screening test especially for early-stage disease.

## Availability of data and materials

All data generated or analyzed during this study, if not included in this article and its supplementary information files, are available from the corresponding author on reasonable request for research use.

## Authors’ contributions

FB, ZW, and Yulong L designed the study and wrote the main manuscript text. Yuanyuan H oversaw next-generation sequencing experiments. TH, FL, SY, SL, XL, and PN performed data analysis and figure preparation. RZ, MZ, PS, FF, WG, JD, GB, Yuan L, QH, and BZ enrolled patients and collected clinical data. Yu H, WC, FT, SG oversaw the study. All authors reviewed and discussed the manuscript.

## Acknowledgements

We thank Shuying He for assistance with exploratory data analysis. The authors are grateful to all patients and their families for their voluntary participation in this study.

