## Supplemental Figures for "Noninvasive cancer detection by extracting and integrating multi-modal data from whole-methylome sequencing of plasma cell-free DNA"

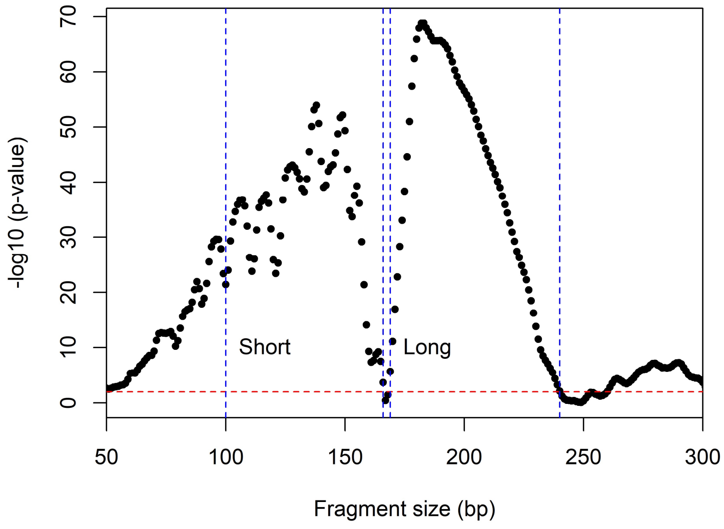


**Figure S1. Difference in cfDNA size distribution between cancer patients and healthy controls.** Wilcoxon test is performed at each fragment size between the frequencies of cancer patients and healthy controls in the training cohort, and –log10 of p values are plotted against fragment sizes. The red dotted line indicates the threshold of significance, p values above which (0.01) are considered insignificant. Blue dotted lines mark the size boundaries of short (100–166 bp) and long (169–240 bp) fragments.

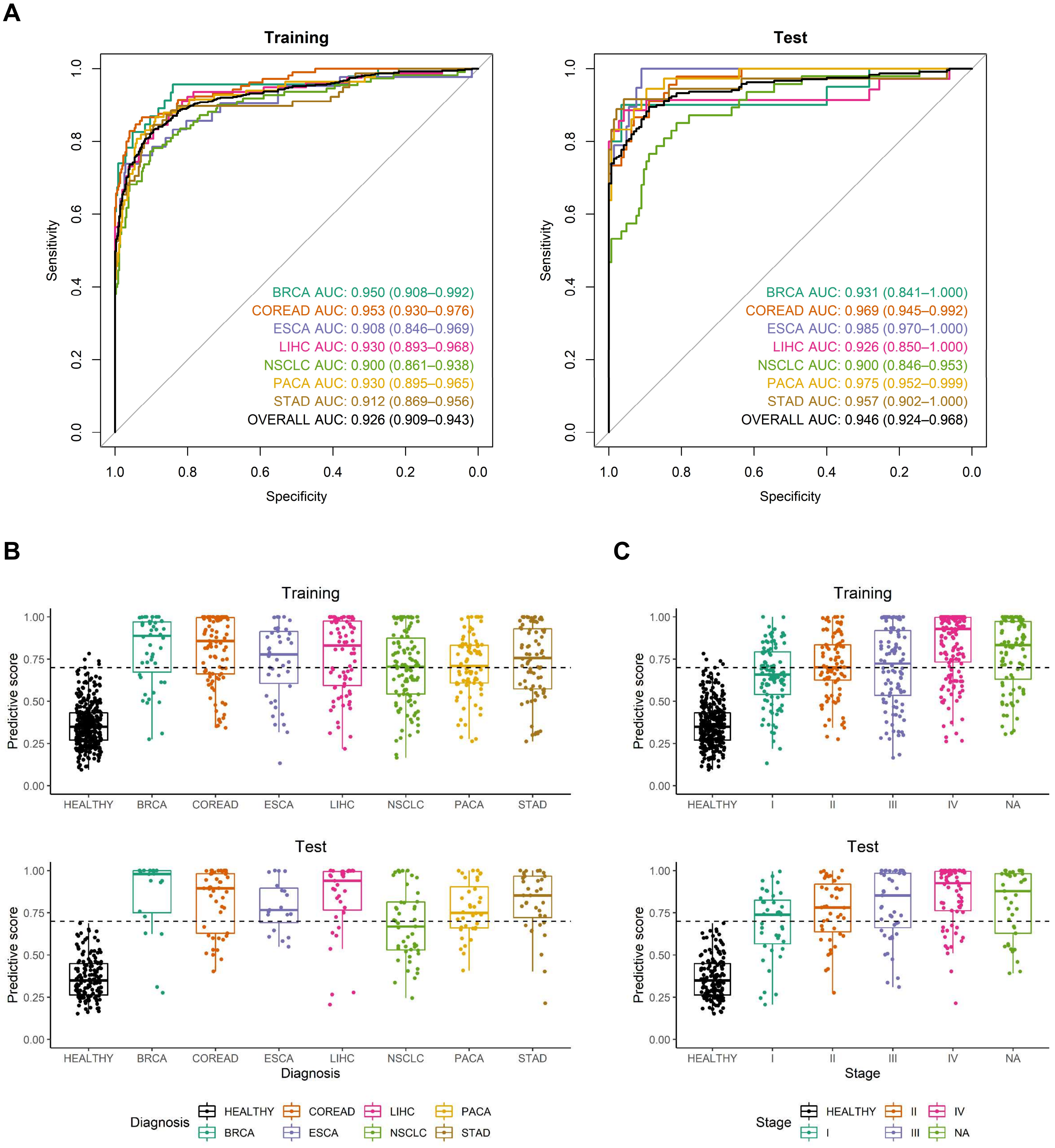


**Figure S2. Cancer detection performance of MFR.** (A) ROC curves for the detection of seven types of cancer using MFR. (B) Predictive scores of MRF by cancer type. (C) Predictive scores of MFR by histological stage. The dotted line indicates the threshold (positive or negative) at 99% training specificity.

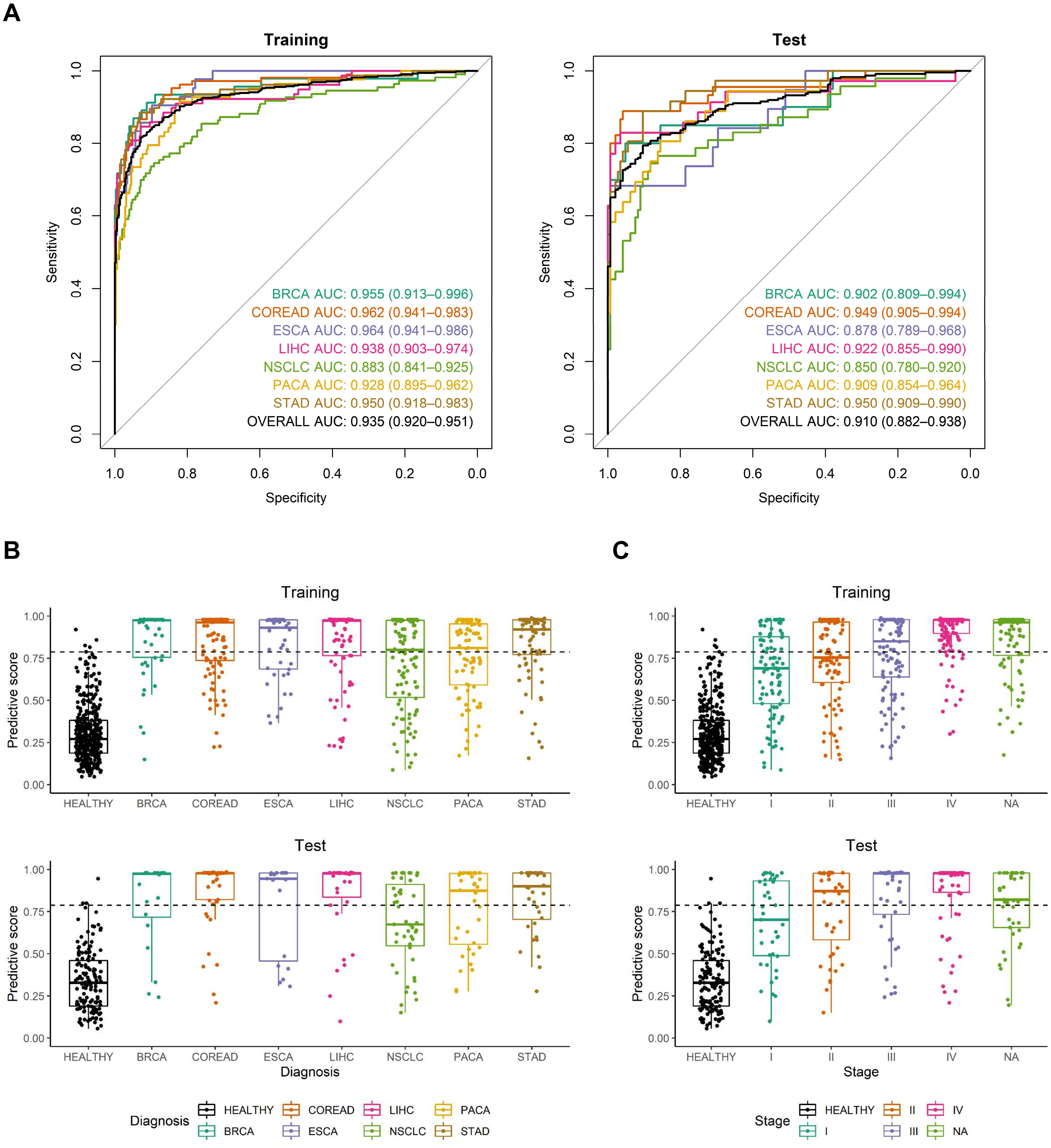


**Figure S3. Cancer detection performance of FSI.** (A) ROC curves for the detection of seven types of cancer using FSI. (B) Predictive scores of FSI by cancer type. (C) Predictive scores of FSI by histological stage. The dotted line indicates the threshold (positive or negative) at 99% training specificity.

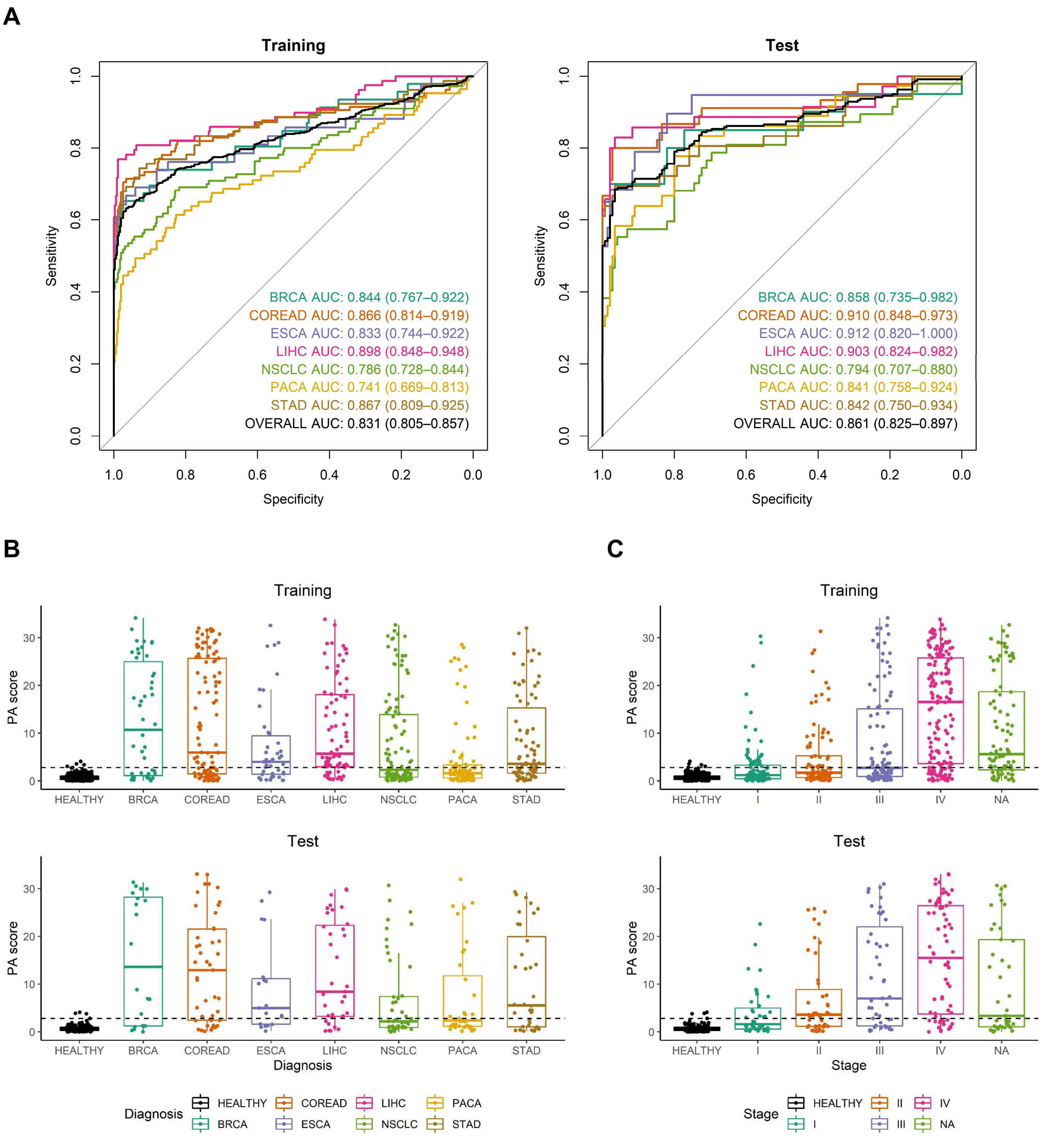


**Figure S4. Cancer detection performance of CAFF.** (A) ROC curves for the detection of seven types of cancer using CAFF. (B) PA scores of CAFF by cancer type. (C) PA scores of CAFF by histological stage. The dotted line indicates the threshold (positive or negative) at 99% training specificity.

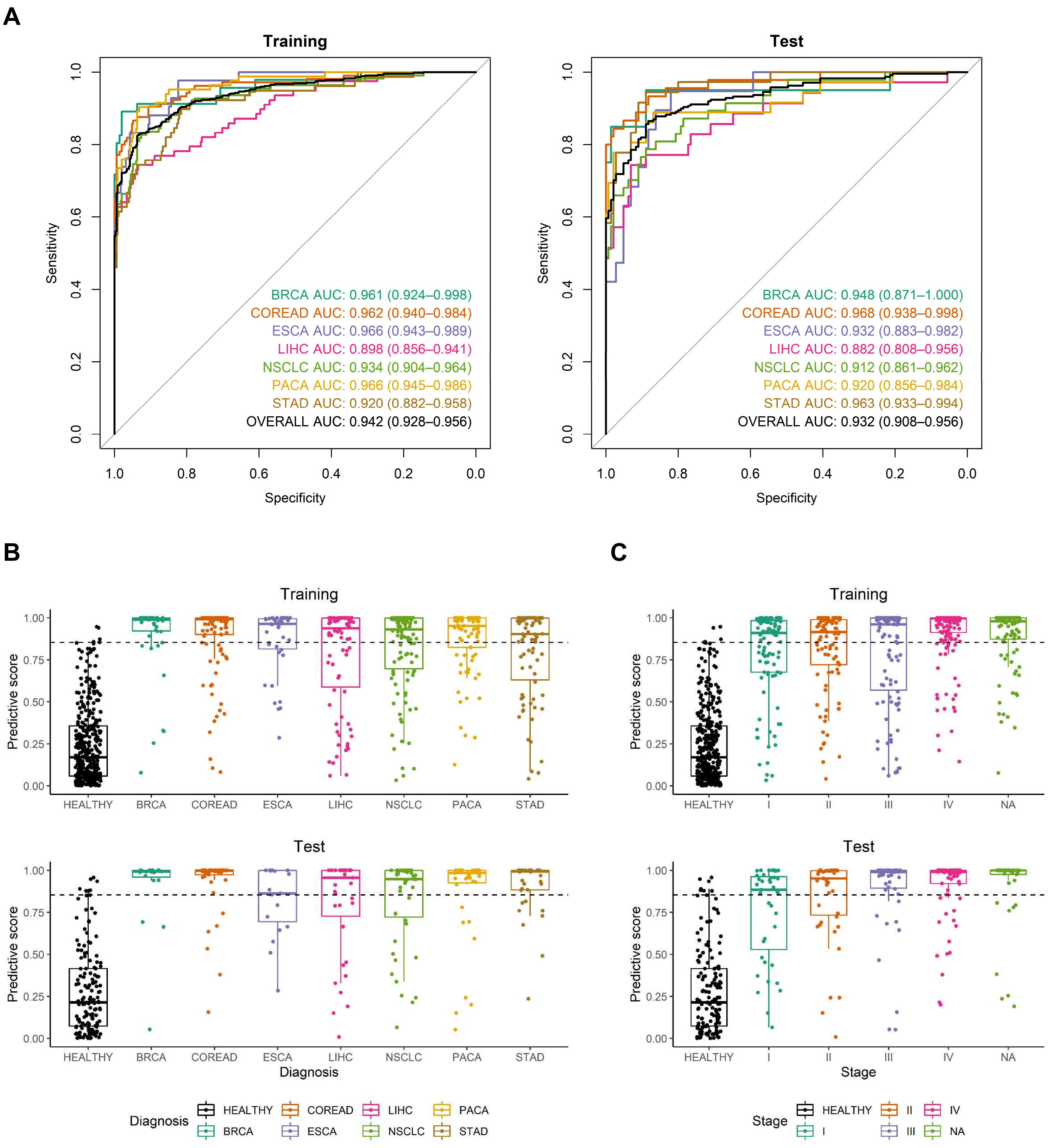


**Figure S5. Cancer detection performance of FEM.** (A) ROC curves for the detection of seven types of cancer using FEM. (B) Predictive scores of MRF by cancer type. (C) Predictive scores of FEM by histological stage. The dotted line indicates the threshold (positive or negative) at 99% training specificity.

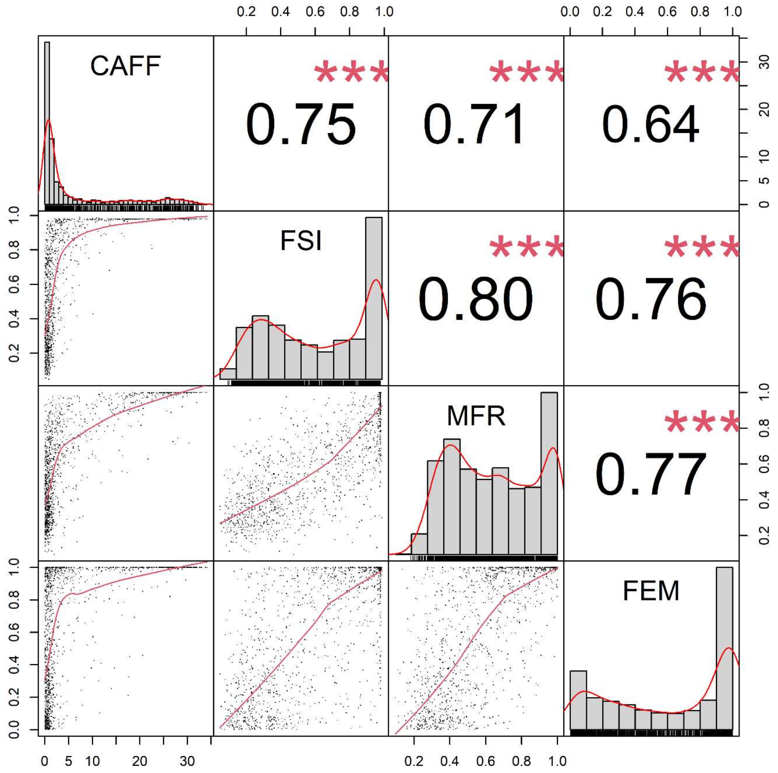


**Figure S6. Correlation between individual biomarkers extracted from WMS data.** Histograms depict the distributions of PA scores of CAFF and predictive scores of FSI, MFR, and FEM respectively. Scatter plots depict the pairwise correlations between biomarkers and regression lines are fitted. Spearman correlation coefficients are calculated and asterisks (***) denote p < 0.001.

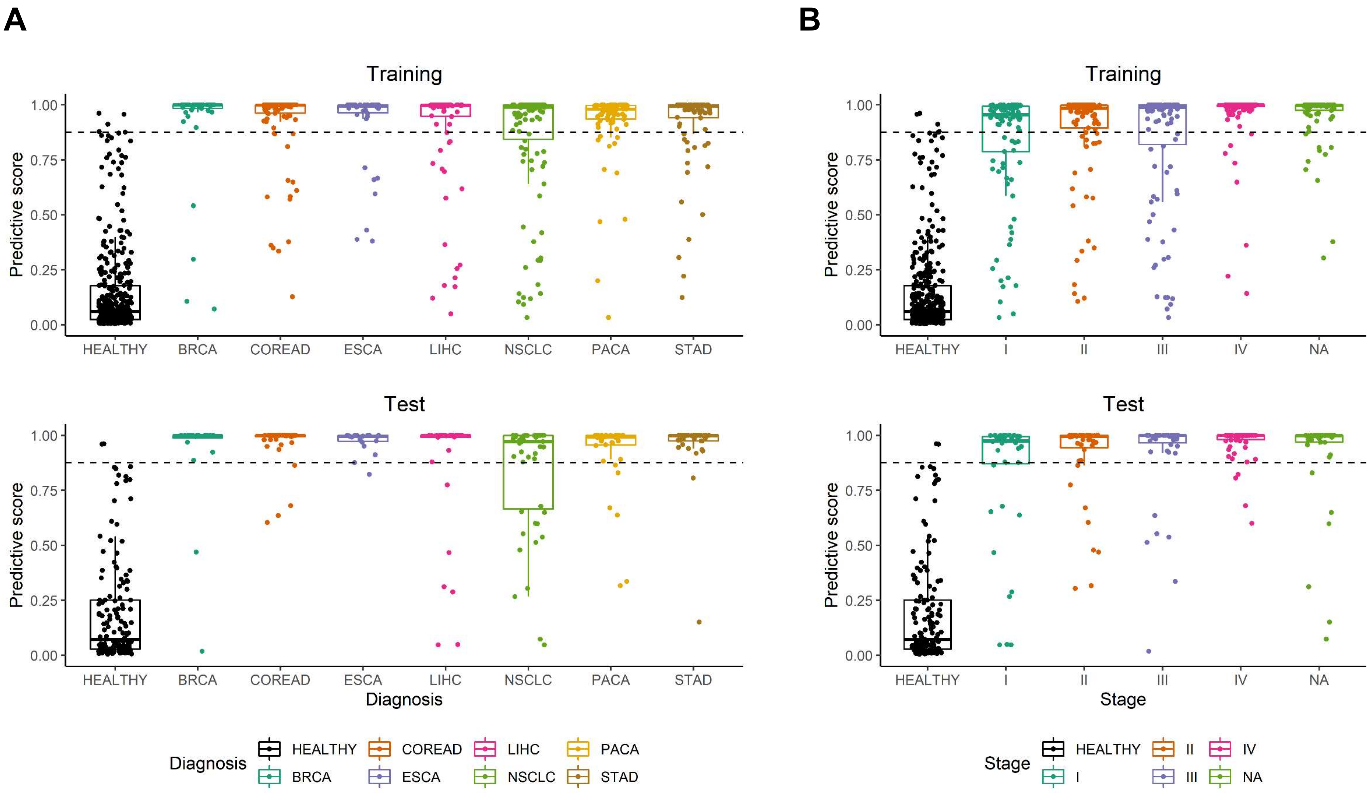


**Figure S7. THEMIS prediction scores by cancer type and histological stage.** (A) Predictive scores of THEMIS by cancer type. (B) Predictive scores of THEMIS by histological stage. The dotted line indicates the threshold (positive or negative) at 99% training specificity.

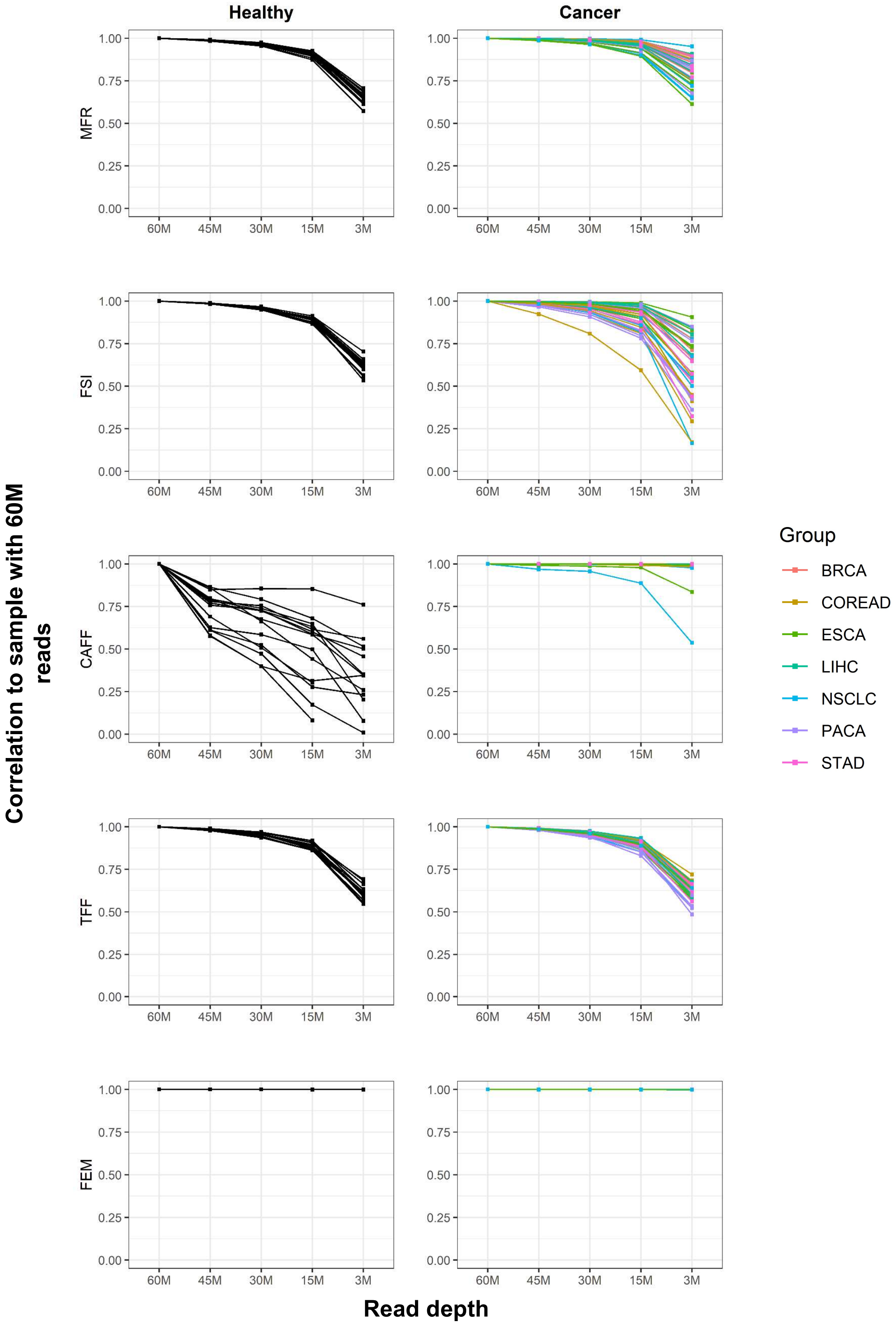


**Figure S8. Comparison of subsampled profiles to initial profile with 60M reads for individual biomarkers.** WMS data from 15 healthy controls and 35 cancer patients (including 5 samples for each cancer type) were randomly subsampled from data with 60M paired reads to 45M, 30M, 15M, and 3M reads respectively. Pearson correlation of feature profiles at each read depth was calculated against data with 60M reads for individual biomarkers. Cancer types are depicted by color.
